# TransgeneR: a one-stop tool for transgene integration and rearrangement discovery using sequencing data

**DOI:** 10.1101/462267

**Authors:** Guofeng Meng

## Abstract

Genetically modified organisms are widely used in lifescience research, agriculture and in commercial products. However, in most cases, the genetic modification in the host genome is often less well characterized with respect to integration location, copy number and host gene expression. The application of next generation sequencing technologies has enabled the characterization of transgene events but still limited by the lack of computational tools. We present a one-stop R tool, transgeneR, as a general computational tool for discovering transgene integration and rearrangement in the host genome. It especially considers the properties of transgene events, such as the homologous transgene sequences, complex genetic structure and multiple copies of transgene insertion. Using this tool, we have successfully mapped the chromosomal transgene integration loci and transgene rearrangements in an artificially simulated MAPT transgene mice genome as well as in a newly generated human tau (MAPT, 0N4R) transgene mice. When unbiased sequencing data such as the whole genome sequencing data, were provided as input, transgeneR integrated multiple information, including integration location, direction, split- and nonsplit-reads, to predict the transgene fragments and their copy number. Overall, our initial evaluation indicates that the transgeneR package is an effective tool for the identification and characterization of transgene integration and rearrangements events, especially in transgene genome with complex genetic structure. TransgeneR is publicly available: https://github.com/menggf/transgeneR

## 1 Introduction

Genetic modification (GM) in domestic species has been a routine technique to study the gene of interest or to generate organisms with the desired phenotypes. GM animals, predominantly, transgene mice are routinely used in biopharmaceutical research to study both the efficacy and safety of molecules prior to testing in humans. This highlights the significance (human safety) and hence the impact (cost) of using GM animals in drug discovery and development [1]. Ironically, attention to the details of the transgene animals are often not taken into consideration while selecting the model systems for pharmacology studies and during the interpretation of results. According to the collection of international mouse strain resource, about 40,000 mice strains have been generated (by Jun. 2017) [2], although how many of them were characterized comprehensively at the genome, gene, transcript, and protein levels remains unclear, especially the transgene generated by pronuclear microinjection that results in random genomic integration. Random integration of a transgene may disrupt an endogenous gene that may either partially or entirely account for the transgene phenotype (e.g. embryonic lethality when the transgene is bred to homozygosity) [3]. In addition, transgenes often integrate as multicopy concatamers resulting in overt expression of transgene product that may influence the phenotype that is entirely artificial. Recent reports for transgene mouse indicate high rate of potentially confounding genetic events [4]. Therefore, a thorough understanding of transgene insertion loci, hemi- and homo-zygosity of the transgene insertion, transgene copy number and its stability in the host genome, at the minimum is essential in the process of generating a transgene animal and interpretation of their phenotypes, and/or while selecting the models for pharmacology studies.

Various methods have been developed to characterize the transgene models, including fluorescence in situ hybridization (FISH) to map the transgene to a chromosome, Southern blotting, restriction mapping/DNA walking, inverse PCR, shot-gun whole genome sequencing, and qRT-PCR [5, 6, 7]. However, each of these methods have their own limitations and no single method can unequivocally address the nature of transgene integration. A combination of methods must be employed to decipher the details. Next generation sequencing has revolutionized genomic/gene expression studies and its utility in GM animals and crops is anticipated to increase. Attempts have been made towards this goal, such as Target Loci Amplification (TLA) [8], VISPA2 [9, 10], CONTRAILS [11] and analysis pipelines [12]. Among them, TLA is an integrated platform by combining experimental targeted locus amplification and sequencing technologies to study the transgene events. Other computational tools usually take advantage of split and discordantly mapped reads to find the host genome break points. They usually have some disadvantages. Most tools do not take into account the complexity of the transgene, including multiple copies of transgene sequences, transgene deletions/rearrangements and integration in the repeat regions. For instance, the single-round reads alignment usually failed to identify the split information in both genome and transgene sequences, which may result in low efficiency usage of split reads and even uncertainty to the predicted integration sites. Not enough consideration to the endogenous homologous sequence of transgene sequence in the host genome also interferes with the interpretation of results, even sometimes result in identification of spurious insertion sites. Meanwhile, most of these tools did not release their codes for a local analysis, which limited their application to the large-size sequencing data.

In this work, we present a new R package, transgeneR, which is designed as a general computational tool to elucidate the transgene integration site and rearrangement using sequencing data e.g. whole genome sequencing (WGS) and amplification-based sequencing data. It applies a two-round alignment and assembles the split reads to predict the transgene integration and rearrangement events. When WGS data are provided as input, transgeneR can predict the transgene fragment usages that results from the rearrangement of transgene sequence. When applied to a simulated transgene mice, TransgeneR successfully predicted all the integration sites and rearrangements while filtering the noise signals. Using the WGS data, transgeneR accurately predicted the transgene fragment, especially the full-length insertion of transgene sequences, and their copies. TransgeneR also predicted both genome integration sites and transgene rearrangements from an experimentally derived WGS data and PCR-based sequencing data of a newly generated transgene mice. Overall, our initial evaluation with both simulated and experimentally derived sequencing data suggests transgeneR is an effective tool to map the transgene information in genetically modified organisms.

## 2 Methods

### 2.1 transgene animals

All animals were bred and handled at GlaxoSmithKline according to the Institutional Animal Care and Use Committee guidelines. Animals were housed on a 12-hour light/12-hour dark cycle (7:00 am to 7:00 pm) in a barrier facility, with controlled temperature and light. The human cDNA of the 0N4R isoform of tau with a single coding variant (P301L) driven by the mouse calcium/calmodulin-dependent protein kinase II alpha (CaMkII*α*) promoter was microinjected into the C57BL/6J embryos and were implanted into surrogate animals. Founders carrying the hTau cDNA and stably transferring the transgene to subsequent generations by germline transmission were selected, and the transgene expression was confirmed by western blot and/or immunohistochemistry, and the selected line (TauD35) was maintained as a colony.

Animal study protocal was reviewed and approved by Institute of Animal Care and Use Committee (IACUC), AUP No. is 0084.

### 2.2 Whole genome- and PCR-based sequencing data

The transgene mice genomic DNA was extracted and purified from the tail to generate the library for sequencing according to the manufacturer’s instructions (Illumina Hiseq 2500 and X10).

A set of primers, the forward primer covering the promoter region at “GGCCTCCCTGTCCATAGA” and the reverse primer of region in Tau “AAGTTCCTCGCCGTCATC”, were synthesized to amplify the integrated cDNA. Mouse tail genomic DNA was extracted by Qiagen kit (Catl#51304). PCR was carried out for 30 cycles at the condition of denature at 95°C for 30sec, and annealing at 58°C for 30sec, elongation at 72°C for 30sec. The amplified DNA was separated in 1% Agarose gel with 1Kb DNA ladder (Invitrogen Catl#10787018). A band around 800 base pairs was expected in the transgene line. Sequencing was performed using the Illumina X10 platform.

### 2.3 Sequencing data analysis

The paired-end sequencing data were subjected to quality control analysis using Fastqc and trimmed the reads with low quality. The duplicated reads were filtered using the methods implemented in ShortReads [13]. The reads were aligned to both mouse genome (mm10) and transgene sequence using bowtie2 [14] under a mode of local alignment. The reads with compensating alignment in the genome and the transgene sequence were proposed as a potential site of integration or rearrangement. The non-aligned clipping parts of the reads were removed in second-round alignment. The alignments were assembled by combining with the first-round alignments.

### 2.4 Homologous sequence

The transgene sequence was artificially fragmented into 200 bp allowing a partial overlap of about 50 bp. Then, all the fragments were aligned to the host genome using bowtie2 and the consensus parts of the transgene sequence were assembled as the homologous parts of the transgene sequence. The homologue information, including the genomic range and its directions were recorded for subsequent analysis. The predicted results were validated by comparison to the transgene sequencing information (http://www.informatics.jax.org/allele/MGI:5646621).

### 2.5 Score of predictions

The alignment scores of split reads are used to indicate the confidence of predicted integration and rearrangement. For a split read *i*, if its alignment score of left and right sites of split reads are 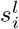 and 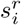, the overall score for this split read is:

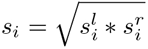

The score of transgene integration or rearrangement site is calculated as the mean value of all the scores of split reads that cross this site

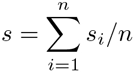

### 2.6 transgene fragment usage

Fragments of transgene sequences may be inserted into the genome either at the same loci where the complete transgene is inserted or at alternate genomic locations. To detect the transgene fragments, transgeneR models the fragment usage based on sequencing depth. In this process, transgeneR estimates the overall sequencing depth using the reads mapped to chromosomes, and then normalize the observed sequencing depth of transgene sequence into copies (*C*). The tool assumes that the *C* values are the sum of all the transgene fragments.

To find the transgene fragments, transgeneR firstly constructs a set of transgene fragment *X* = *{x*_1_*, x*_2_*, .., x_i_}* by connecting the starting (S) and ending (E) points in transgene integration and rearrangement events. TransgeneR applies a non-negative linear model implemented in nnls package to fit for the *C* values:

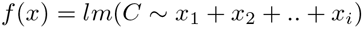

To find integer copies of transgene fragments, transgeneR selects the fragments with coefficents around 1 or other integer values and makes them the initial sets of transgene fragments. Other fragments are added or removed to check if it can improve the correlation value *r* of the fragment copies (*Y*) and the sequencing depth (*C*), where

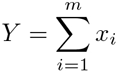
and *m* is the number of selected fragments.

## 3 Results

### 3.1 Overview of TransgeneR

TransgeneR is designed to identify the transgene integration and rearrangement events mainly by capturing the split and discordant mapped reads. Given that more often than not a transgene model will carry a homologous sequence in the host genome, transgeneR annotates the homologous regions based on sequence similarity and this information can be used to determine the origin of reads when they are mapped to such regions.

A schematic pipeline of the transgeneR tool is described in Figure 1(a). Briefly, the paired-end reads are mapped to both the host genome and the transgene sequence using bowtie2 with the same parameter setting [14]. To identify the split reads, soft clipping is allowed by applying local alignments. When the whole genome sequencing data is provided as input, most of reads are originate from the genome sequences, and therefore, the reads with the exact mapping to the genome are filtered from further considering.

**Figure 1:**
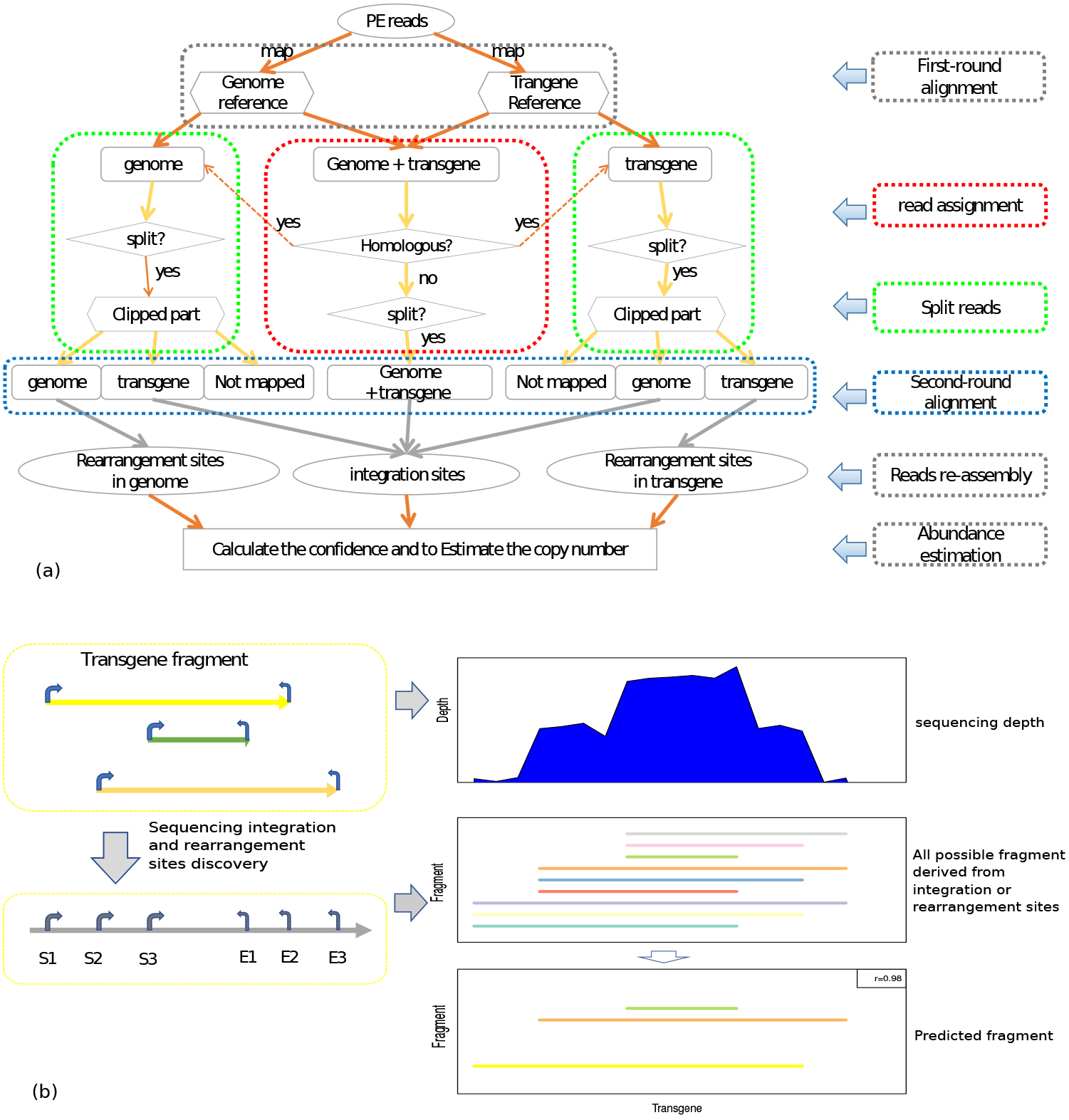
The workflow of transgeneR. (a) The pipeline for transgene integration and rearrangement site discovery. (b) The method to predict the transgene fragment usage.

Next, transgeneR performs the first-round reads assignment for the reads mapped to both the genome and transgene sequence. If one read is split mapped into both genome and transgene sequences, e.g. first half mapped to the genome and the second half mapped to the transgene sequence, it will be considered as evidence of integration and therefore will add a count of 1 to this location. If the mapped part of one read overlaps, i.e. the first half of the read is mapped both in the genome and the transgene sequences, this read will be uniquely assigned to either the genome or the transgene sequence according to the mapping score, mapping concordance and homologue annotation. Since the probability of random integration of the transgene into the homologous gene loci in the host genome is extremely rare or negligible, reads mapped to the homologous regions are preferentially assigned to the integration sites if equal mapping scores are observed.

Then, soft-clipping part of partially mapped reads are cut as fragments and re-aligned to both the genome and transgene sequences. Second round of read assignment is performed to ensure each fragment has a unique mapping location. Each mapped fragment will be counted as one evidence of split site from the original mapped location to the fragment mapped location. In this step, new mapping location can be at different sequences, e.g. the location of original mapping is on the host genome while the fragment is mapped to the transgene sequences. Therefore, the scenario can be reported: (a) transgene integration, where part of the mapped reads is on chromosome and the remaining in transgene sequence; (b) genomic rearrangement, where two parts of the reads are both mapped to the chromosomes; (c) transgene rearrangement, where two parts of the reads are both mapped to the transgene sequence. Finally, the reads with the same split location are summarized for the prediction of transgene integration and rearrangement sites. In this process, the number of split reads is the most critical criteria. Only the sites with enough split reads are considered for the final prediction. TransgeneR also takes into consideration other factors, such as the balanced distribution of mapped reads around predicted sites, the read split patterns, complexity of predicted genomic location and quality of mapping. In the final output, transgeneR reports multiple information for each integration or rearrangement site, including (a) the number of split, nonsplit and crossing reads; (b) organization directions, such as forward-forward (ff), forward-reverse(fr), reverse-forward (rf) and reverse-reverse (rr); (c) fragment gaps; (d) paired host genome locations of integration sites. TransgeneR also generates the visualization of read alignment around the discovered sites (see Figure 3(a)).

When non-biased sequencing data are used, TransgeneR can predict the inserted transgene fragment and their copies (see Figure 1(b)). The integration and rearrangement sites are firstly assigned as either starting (S) or ending (E) points based on their organization orientation in the host genome. All the possible transgene fragments are generated by connecting S and E points. TransgeneR applies a method to find the subset combination of transgene fragments so that the estimated fragment coverage has the best correlation with the sequencing depth. In this step, this ratio of split and nons-split reads can be used to estimate the usage frequency of S and E points, which determines the copies of fragments derived from the same S or E points.

### 3.2 Evaluation with simulated transgene genome

To evaluate the performance of transgeneR, we generated an artificial transgene mice genome using Tg(Camk2a-MAPT*P301L)D3 () as the transgene sequence. We arbitrarily designed the transgene genome to have three transgene integration events (chr1, chr6 and chr17), and a total of 12 transgene rearrangement fragments, with 6 integration sites (1-6) and 13 rearrangement sites (a-k) in the mouse genome (see Figure 2(a)). Among them, rearrangement sites (a), (c) and (e) occurred twice. The simulated WGS data was generated by ART [15] with the setting of (1) 50X read depth, (2) HiSeq 2500 sequencing platform, and (3) 150 bp paired-end reads.

**Figure 2:**
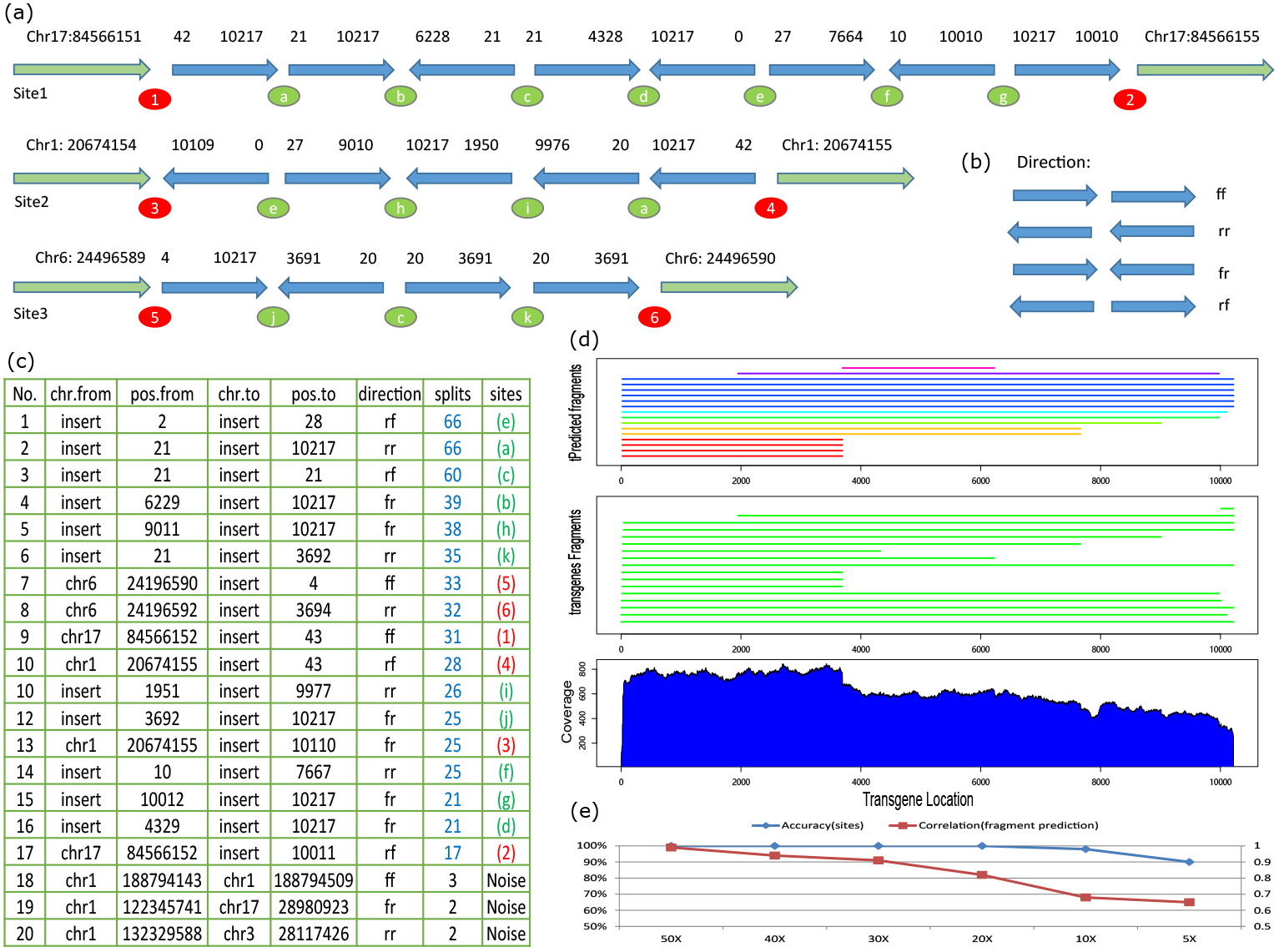
Computational evaluation using simulated whole genome sequencing data. (a) the simulated mice transgene genome, including three transgene integration on chr17, chr1 and chr6; (b) the definition of transgene sequence organization direction using 4 two-letter phrases; (c) predicted the transgene integration and rearrangement sites, including the genomic/transgene locations (chr.from, pos.from, chr.to and pos.to), direction (direction) and number of supporting reads (splits); (d) the predicted transgene fragment usage.

Analysis using transgeneR showed that it successfully discovered all the integration sites (a-k) and rearrangement sites (1-6) (see Figure 2(c)). Number of split reads was used as a measure of confidence on integration or rearrangement sites. While the median score for all the true integration sites was 31, the minimum score was 17, which was observed with the transgene integration site (2) on the left side of chromosome 17. In this artificial genome, the (a), (c) and (e) have two rearrangement sites. Consequently, about twice the number of the median split reads were observed. Besides true rearrangement events or integration sites, transgeneR also reported other false-positive sites. However, they usually had significantly less split reads to support them, e.g. only 3 split reads, which is far less than the minimum number of true sites. Overall, it is evident that transgeneR can detect the true transgene integration and rearrangement sites, and that the number of rearrangement sites appears to be related linearly with the split read number. In addition, transgeneR also recovered the orientation of all the transgene fragments in the insert sequences (see Figure 2(b, c)).

Moreover, transgeneR estimated the transgene fragment length and copy numbers in the transgene genome while using the WGS data. In this model, the coverage calculated from WGS data are modeled as the sum values of transgene fragment, which is initialized by the continuous combination of integration or rearrangement sites. Using the simulated genome, the predicted coverage by summing up the predicted fragments and their copies, has a Pearson’s correlation of *r* = 0.995 with the true coverage values. Figure 2(d) shows the predicted results for transgene fragments. Comparing with the true fragments of the simulated transgene genome, transgeneR recovered 90% of transgene fragments and 100% of the complete transgene sequences.

To evaluate the robustness of transgeneR, we performed another round of evaluation by decreasing the sequencing depth from 50X to 5X. The accuracy of integration and rearrangement site discovery is displayed in Figure 2(e). Overall, transgeneR could still discover the true integration and rearrangement sites. However, when the sequencing depth decreased, the ability of transgeneR to discriminate the false sites decreased accordingly. We also predicted the fragment usage. Figure 2(e) showed the correlation values between the estimated sequencing depth using predicted transgene fragments and the true sequencing depth, which suggested that prediction accuracy for transgene fragment usage was more sensitive to sequencing depth.

Together, these results indicate that transgeneR not only accurately predicted the transgene integration and rearrangement sites but also transgene fragment usage, especially the complete insertions.

### 3.3 Evaluation of transgeneR using experimentally derived whole genome sequencing data

A human MAPT (0N4R) cDNA containing the P301L variant driven by the CaMKII*α* promoter was cloned and was microinjected into embryos to generate the human tau transgene mice. The genotype of the founder lines were evaluated using transgene specific PCR to determine successful germline transmission as well as by qPCR to determine relative levels of the expression of the transgene. One such line in the C57Bl6/J background, named herein after as TauD35, maintained as a heterozygote, was subjected to whole genome sequencing analysis by HiSeq X10.

Following quality control and filtering the duplicate reads, transgeneR was applied for the analysis of transgene integration and rearrangement. Actual sequencing depth was estimated by analysis to chromosome 1 and the average depth was determined to be 49.5X. Considering that the transgene mice is a heterozygote, the expected split reads based on this coverage is not more than 25. Unlike the artificially simulated WGS data, analysis of the experimentally derived WGS data will have certain challenges, especially in the repeat regions. Because of this, it is impossible to use number of split reads as the only criteria to select the transgene integration sites. Therefore, transgeneR also considers whether the split reads show a balanced reads distribution across the predicted sites i.e. integration sites with clipping fragments mapped only to one side will not be selected as a true integration site. Our evaluation works also found that it was an effective way to filter the false prediction by considering the DNA sequence complexity of mapped genomic regions. TransgeneR evaluates the genomic complexity by checking the uniqueness of the mapped reads. The split reads mapped to the same type of complex region, e.g LINE, are filtered as integration or rearrangement site discovery. All these strategies help to identify true integration sites. Analysis of the sequencing data suggested a single transgene integration site on chromosome 4 [chr4:129387127 (left side) and chr4:129387416 (right side)] (see Figure 3(a)). It was supported by 19 split reads and 18 split reads to the left and right sides, respectively. In addition, there were also 8 reads cross this integration site.

**Figure 3:**
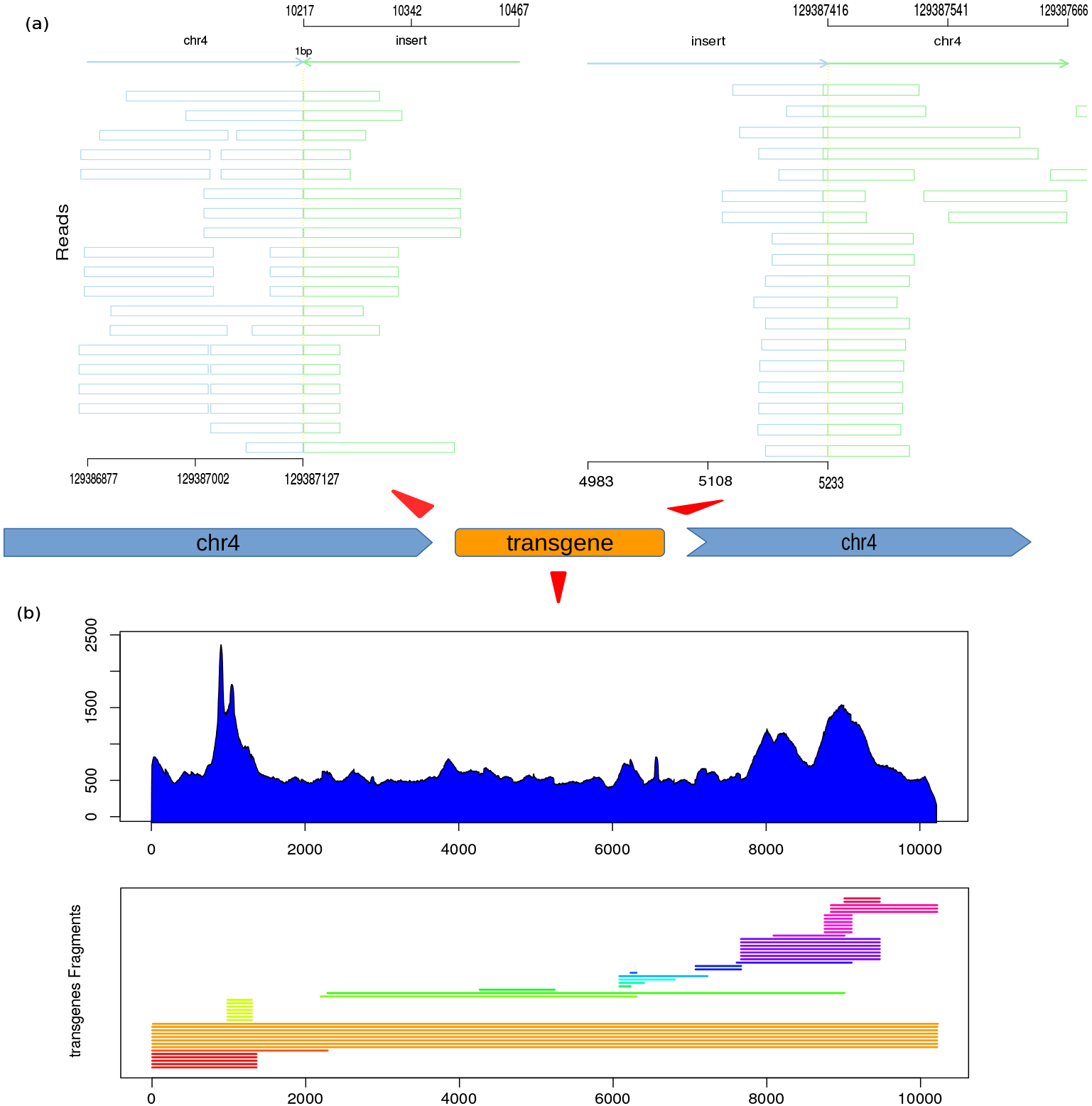
Application of transgeneR to the WGS data. (a) The predicted transgene integration sites (including the left side and right site) and visualization of the split reads; (b) the predicted transgene fragment usage.

Furthermore, transgeneR also identified about 20 transgene rearrangement sites in the genome, majority of which were in the head-tail orientation of complete transgene sequences. By counting the reads mapped to the transgene sequence, it was equivalent to about 16 copies of the transgene sequence. TransgeneR predicted the fragments that were resulted from the transgene rearrangements (see Figure 3(b)). Among them, there were 8 full copies and others were partial copies. Along the sequencing depth plot, there were two regions, around 1200 and 9500 bp, that had higher read coverage. The technical reason for this is unclear at present. They seem to be the hotspot regions with transgene rearrangement. TransgeneR predicted many short fragments in these two regions.

### 3.4 Utility of transgeneR to PCR-based sequencing data

PCR-based sequencing methods may also be used to map the transgene insertion loci using the transgene specific primers and degenerate primers that can anneal to the host genome. Such a method was also employed in the tail DNA isolated from TauD35 mice, sequenced using MiSeq, and analysed using targetR, to evaluate the compatibility of this package with sequences obtained using different methodologies.

Unlike WGS data, most of the split reads (*>* 95%) mapped to the transgene rearrangement sites, especially the head-tail rearrangement of complete transgene sequences. This is expected, and reflects targeted enrichment of the transgene since sequence specific primers were used in combination with degenerate primers. Table 1 lists the predicted rearrangement sites by transgeneR and most of them were found to be consistent with the results from WGS analysis. Interestingly, besides the id entification of rearrangement sites, the integration site on chromosome 4 is also observed, albeit only by one split read, although the reason for this is currently unclear.

**Table 1:** Predicted integration and rearrangement sites using WGS and PCR-based sequencing data

| PCR-amplification |  |  |  |  |  | WGS |  |  |  |  |  |
| --- | --- | --- | --- | --- | --- | --- | --- | --- | --- | --- | --- |
| chr.from | pos.from | chr.to | pos.to | dir | reads | chr.from | pos.from | chr.to | pos.to | dir | reads |
| transgene | 2 | transgene | 10217 | rr | 1739 | transgene | 3 | transgene | 10217 | rr | 1556 |
| transgene | 4 | transgene | 4328 | rf | 1726 | transgene | 4 | transgene | 4325 | rf | 153 |
| transgene | 2 | transgene | 7664 | rr | 1483 | transgene | 3 | transgene | 7666 | rr | 147 |
| transgene | 2 | transgene | 9010 | rr | 1403 | transgene | 3 | transgene | 9010 | rr | 149 |
| transgene | 4 | transgene | 6228 | rf | 1124 | transgene | 4 | transgene | 6228 | rf | 159 |
| transgene | 4 | transgene | 2223 | rf | 1070 |  |  |  |  |  |  |
| transgene | 4 | transgene | 1950 | rf | 633 | transgene | 4 | transgene | 1950 | rf | 154 |
| transgene | 4 | transgene | 93 | rf | 304 | transgene | 4 | transgene | 74 | rf | 149 |
| transgene | 52 | transgene | 10106 | rf | 170 |  |  |  |  |  |  |
| transgene | 6402 | transgene | 7075 | ff | 118 | transgene | 6402 | transgene | 7075 | ff | 149 |
| transgene | 4 | transgene | 70 | rf | 97 | transgene | 4 | transgene | 74 | rf | 149 |
| transgene | 152 | transgene | 8896 | rf | 87 |  |  |  |  |  |  |
| transgene | 4 | transgene | 128 | rf | 86 |  |  |  |  |  |  |
| transgene | 152 | transgene | 8912 | rf | 81 |  |  |  |  |  |  |
| transgene | 4 | transgene | 10 | rf | 77 |  |  |  |  |  |  |
| chr4 | 129387127 | transgene | 10217 | fr | 19 | chr4 | 129387128 | transgene | 10217 | fr | 1 |

### 3.5 Comparison with independent tools

Currently, limited tools has been specifically designed for transgene integration and rearrangement studies or release their codes. We compared the performance of transgeneR with VISPA2 [10]. Using the same

### 3.6 Performance

TransgeneR tool provides the option to use flexible computational power. Multiple-threads computation is supported for nearly whole analysis process, especially the heavy-computational parts. Users can set the “cores” parameter to define how many threads to use and “buffer size” option to adjust the memory usage. Less the “buffer size”, less memory will be required. TransgeneR was tested for its performance on a 4-core (8 threads) AMD Ryzen 1400 CPU and 16G ROM computer in this study. When 6 threads were used, the average running time for 50X WGS data of the simulated transgene mice genome was about 11.5 hours (not including the time for bowtie2 reads alignment) and the maximum memory usage was about 12GB at a “buffer size” setting of 500,000. The tool was also evaluated using the PCR-amplification based sequencing data that had about 0.5 − 2 million reads. The average analysis time was less than 6 minutes. During the analysis, many temporary files will be generated as output and this may require a large disk space. Users can also set to compress the input .fastq files and temporary files, which may save about 70% memory storage.

## 4 Discussion

TransgeneR is designed as an integrated tool for the discovery of transgene integration site and rearrangement events. Different from the existing tools, it is developed as a general tool fit for sequencing data generated by different methods i.e. whole genome sequencing and amplification-based sequencing. For ease of use, the entire package has only one R function as the centralized interface, even though the whole analysis is divided into multiple steps. In case that the users encounter errors or need to adjust parameters, transgeneR supports the analysis to only part of the whole pipeline by outputting the analysis results of each step into readable files. TransgeneR can automatically judge the output of each steps and decides which steps have been performed. Users can re-run part of the analysis just by deleting the corresponding output files (see package vignette doc).

TransgeneR uses bowtie2 as the alignment tool and takes paired-end fastq files as input. The whole analysis pipeline is optimized based on the output of bowtie2. Currently, it has not been tested for its compatibility with other alignment tools. To use transgeneR, users should install bowtie2 and make the bowtie2 accessible from system path. The genome reference built using “bowtie2-build” is mandatory for transgeneR package. More alignment tools will be tested, and if compatible will be supported in future versions.

Our evaluation supported transgeneR to have a good performance to recover all the transgene integration and rearrangement sites in both simulated and the experimentally derived genome sequences. TransgeneR has been optimized to output the final results based on a set of criteria, including the number of split reads, reads distribution around the predicted sites and the DNA complexity of predicted regions. However, the complex situations of transgene events may lead to false predictions. For example, the biased distribution of the reads in the low complex regions leads to the non-existence of unique threshold to determine the true transgene integration sites. In many cases, manual evaluation is necessary for a reliable conclusion. In support, TransgeneR output provides some necessary information, including the sequencing depth and genomics visualization of the split reads. Although transgeneR tool is easy to use and its application to the newly generated TauD35 mice identified the hMAPT insertion loci in Chr4 by both WGS and PCR based methods, there are some limitations: (a) the predicted sites were not experimentally validated by alternate methods (e.g. amplification of the entire genome insertion site using sequence specific primers spanning the split reads followed by NGS to both confirm as well as to determine the number of functional copies of the transgene; and (b) experimental validation of the expression levels of the host gene where the transgene was integrated and its influence on whole genome expression. Future studies will address these questions.

This package is initially designed for characterizing genome modifications in transgene animals. However, its design strategy is applicable for other DNA integration related discovery. One example is to predict the virus integration sites in the host genome. This can be achieved by modifying few parameters to adjust the tolerance of bowtie2 to the mismatches in the viral genome.

## 5 Acknowledgements

This work is supported by GSK R&D Shanghai. We thank Michelle Zhu for experimental supports in sample collection.

